# Community structure follows simple assembly rules in microbial microcosms

**DOI:** 10.1101/067926

**Authors:** Jonathan Friedman, Logan M. Higgins, Jeff Gore

**Author notes:** The authors declare no competing financial interests. Correspondence and requests for materials should be addressed to J.F. or J.G.

## Abstract

Microbes typically form diverse communities of interacting species, whose activities have tremendous impact on the plants, animals, and humans they associate with^1–3^, as well as on the biogeochemistry of the entire planet^4^. The ability to predict the structure of these complex communities is crucial to understanding, managing, and utilizing them^5^. Here, we propose a simple, qualitative assembly rule that predicts community structure from the outcomes of competitions between small sets of species, and experimentally assess its predictive power using synthetic microbial communities. The rule's accuracy was evaluated by competing combinations of up to eight soil bacterial species, and comparing the experimentally observed outcomes to the predicted ones. Nearly all competitions resulted in a unique, stable community, whose composition was independent of the initial species fractions. Survival in three-species competitions was predicted by the pairwise outcomes with an accuracy of ~90%. Obtaining a similar level of accuracy in competitions between sets of seven or all eight species required incorporating additional information regarding the outcomes of the three-species competitions. Our results demonstrate experimentally the ability of a simple bottom-up approach to predict community structure. Such an approach is key for anticipating the response of communities to changing environments, designing interventions to steer existing communities to more desirable states, and, ultimately, rationally designing communities *de* novo^6,7^.

## Main text

Modeling and predicting microbial community structure is often pursued using bottom-up approaches that assume that species interact in a pairwise manner^8–11^. However, pair interactions may be modulated by the presence of additional species^12,13^, an effect that can significantly alter community structure^14^ and may be common in microbial communities^15^. While it has been shown that such models can provide a reasonable fit to sequencing data of intestinal microbiomes^16,17^, their predictive power remains uncertain, as it has rarely been directly tested experimentally (^18,19^ are notable exceptions).

Current approaches to modeling microbial communities commonly employ a specific parametric model, such as the generalized Lotka-Volterra (gLV) model^20–22^. Generating predictions from such models requires fitting a large number of parameter values from empirical data, which is often challenging and prone to over-fitting. In addition, the exact form of the interactions needs to be assumed, and a failure of the model can reflect a misspecification of the type of pairwise interaction, rather than the presence of higher-order interactions^23^.

Here we take an alternative approach in which qualitative information regarding the survival of species in competitions between small sets of species (e.g., pairwise competitions) is used to predict survival in more diverse multispecies competitions (Fig. 1). While this approach forgoes the ability to predict exact species abundances, it does not require specifying and parameterizing the exact form of interactions. Therefore, it is robust to model misspecification, and requires only survival data, which can be more readily obtained than exact parameter values.

**Figure 1.**
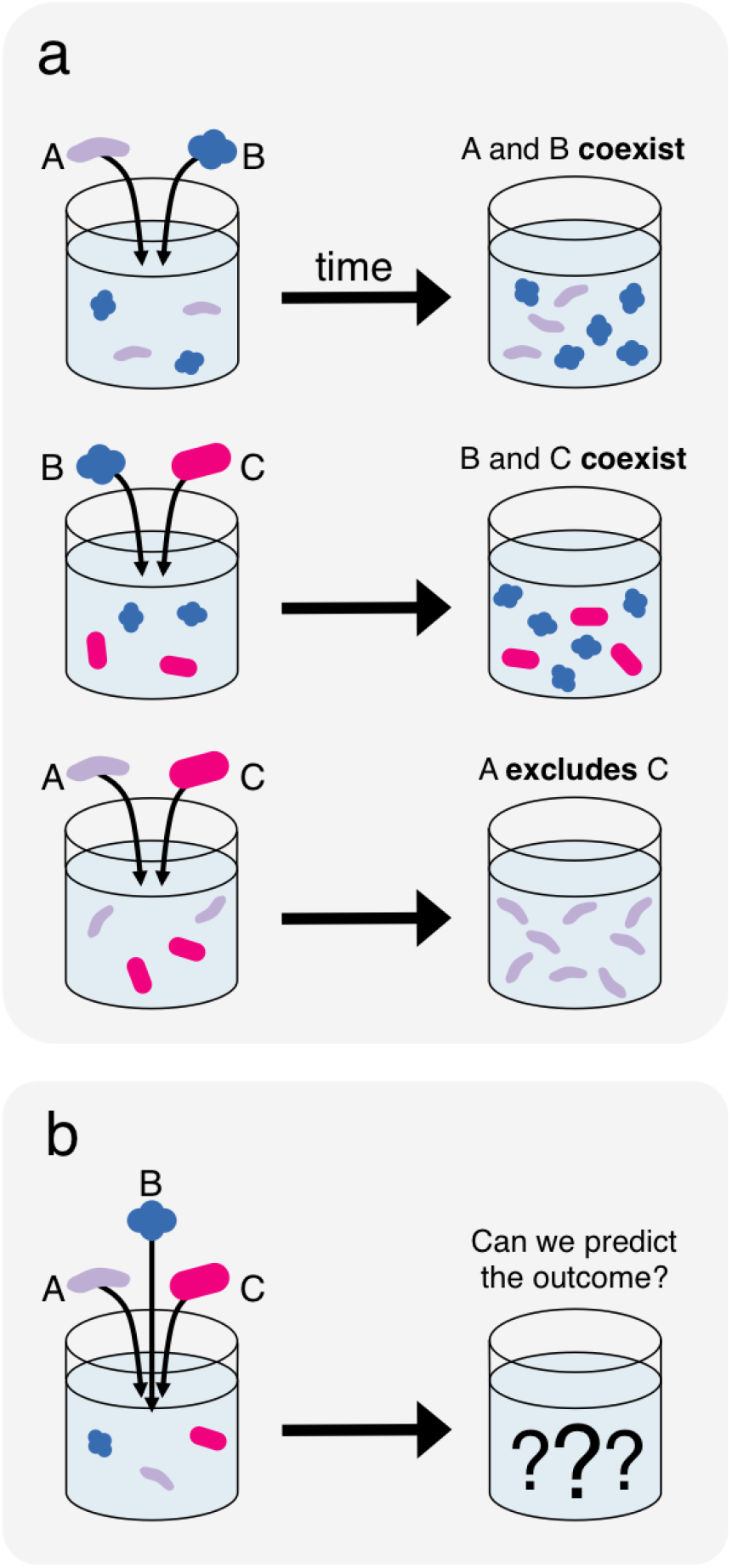
A bottom-up approach to predicting community composition from qualitative competitive outcomes. Qualitative information regarding the survival of species in competitions between small sets of species, such as pairwise competitions (a) is used to predict survival in more diverse multispecies competitions, such as trio competitions (b). The particular pairwise outcomes illustrated here reflect the true outcomes observed experimentally in one set of three species (Fig. 3b).

Intuitively, competitions typically result in the survival of a set of coexisting species, which cannot be invaded by any of the species that went extinct during the competition. To identify sets of species that are expected to coexist and exclude additional species, we first use the outcomes of pairwise competitions. We propose the following assembly rule: in a multispecies competition, species that all coexist with each other in pairs will survive, whereas species which are excluded by any of the surviving species will go extinct. This rule provides a formalizes out intuition, and can be used to systemically predict community structure from pairwise outcomes (Methods, **Fig. S1**).

To directly assess the predictive power of this approach, we used a set of eight heterotrophic soildwelling bacterial species as a model system (Fig. 2a, **Methods**). Competition experiments were performed by co-inoculating species at varying initial fractions, and propagating them through five growth-dilution cycles (**Fig. S2**). During each cycle, cells were cultured for 48 hours and then diluted by a factor of 1500 into fresh media, which corresponds to ~10.6 cellular divisions per growth cycle, and ~53 cellular divisions over the entire competition period. The overall competition time was chosen such that species extinctions would have sufficient time to occur, while new mutants would typically not have time to arise and spread. Community compositions were assessed by measuring the culture optical density (OD), as well as by plating on solid agar media and counting colonies, which are distinct for each species^25^. These two measurements quantify the overall abundance of microbes in the community, and the relative abundances of individual species, respectively. All experiments were done in duplicate.

**Figure 2.**
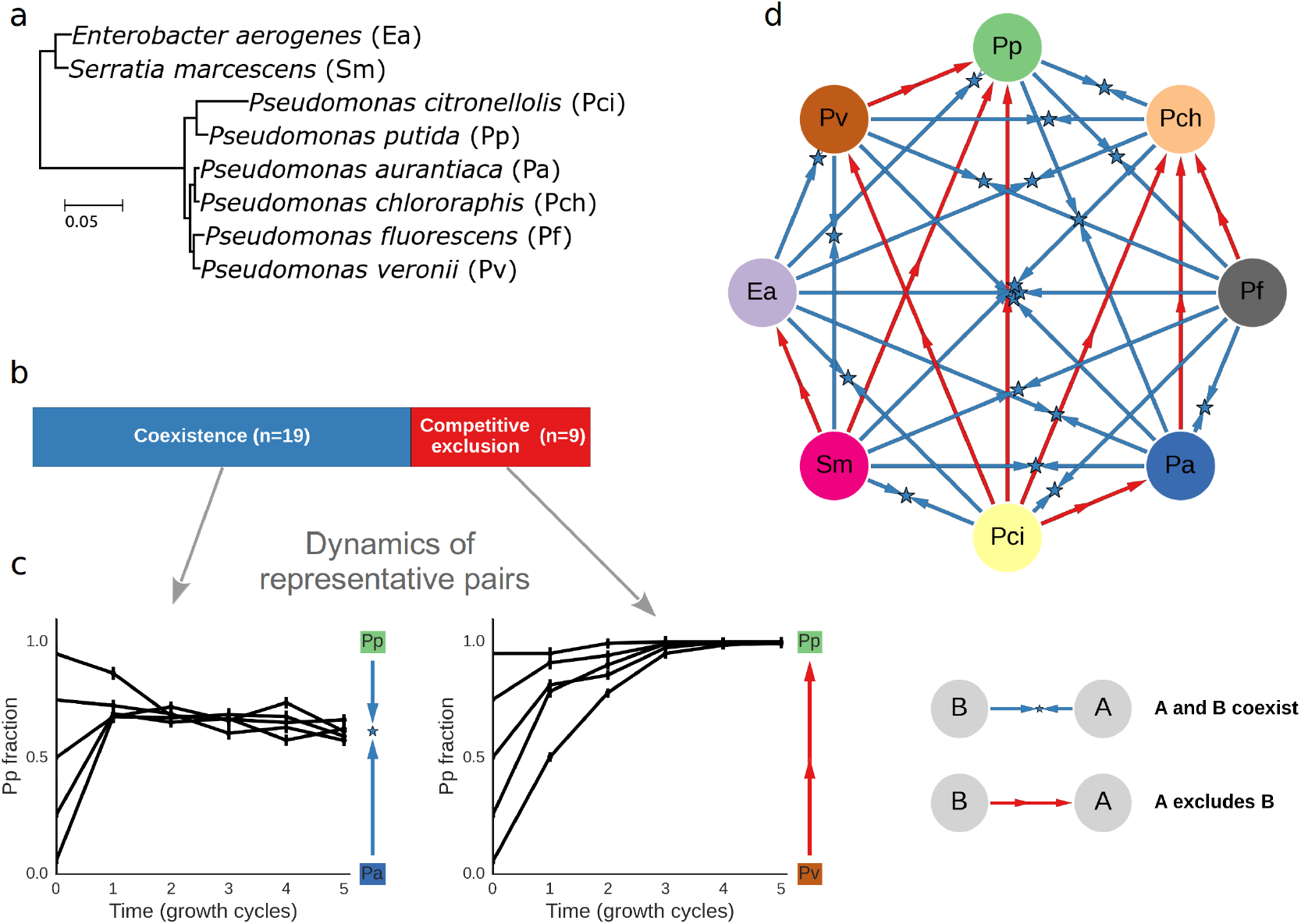
Pairwise competitions resulted in stable coexistence or competitive exclusion. (a) Phylogenetic tree of the set of eight species used in this study. The tree is based on the full 16S gene, and the branch lengths indicate the number of substitutions per base pair. (b) Coexistence was observed for 19 of the 28 pairs, whereas competitive exclusion was observed for 9 of the 28 pairs. (c) Changes in relative abundance over time in one pair where competitive exclusion occurred, and one coexisting pair. The y-axis indicates the fraction of one of the competing species. In the exclusion example, the species fraction increased for all initial conditions, resulting in the exclusion of the competitor. In contrast, in the coexistence case, fractions converged to an intermediate value and both species were found at the end of the competition. Blue and red arrows to the right indicate the qualitative competitive outcome, with the star marking the final fraction in the case of coexistence. Error bars represent the standard deviation of the posterior Beta distribution of the fractions, based on colony counts averaged across replicates. (d) Network diagram of the outcomes of all pairwise competitions.

Pairwise competitions resulted in stable coexistence or competitive exclusion of one of the species. We performed competitions between all species pairs and found that in the majority of the pairs (19/28 = 68%, Fig. 2b) both species could invade each other, and thus stably coexisted. In the remaining pairs (9/28 = 32%) competitive exclusion occurred, where only one species could invade the other (Time trajectories from one coexisting pair and one pair where exclusion occurs are shown in Fig. 2c. Outcomes for all pairs are shown in Fig. 2d). Species' growth rate in monoculture was correlated with their average competitive ability, but, in line with previous reports^26^, it could not predict well the outcome of specific pair competitions (**Fig. S3**).

Next, we measured the outcome of competition between all 56 three-species combinations. These competitions typically resulted in a stable community whose composition was independent of the starting fractions (**Table S1**). However, 2 of the 56 trios displayed inconsistent results with high variability between replicates. This variability likely resulted from rapid evolutionary changes that occurred during the competition (**Fig. S4**). All but one of the other trio competitions resulted in stable communities with a single outcome, independent of starting conditions. This raises the question of whether this unique outcome could be predicted based upon the experimentally observed outcomes of the pairwise competitions.

Trios were grouped by the topology of their pairwise outcome network, which was used to predict their competitive outcomes. The most common topology involved two coexisting pairs, and a pair where competitive exclusion occurs (30/56 = 54%). To illustrate this scenario, consider a set of three species, labeled A, B, and C, where species A and C coexist with B in pairwise competitions, whereas C is excluded when competing with A. In this case, our proposed assembly rule predicts that the trio competition will result in the survival of species A and B, and exclusion of C (Fig. 3a). This predicted outcome occurred for a majority of the experimentally observed trios (Fig. 3b), but some trio competitions resulted in less intuitive outcomes (Fig. 3c). For example, 1 of the 30 trios with this topology led to the extinction of A and the coexistence of B and C (Fig. 3c). The experimentally observed outcomes of competition in this trio topology highlights that our simple assembly rule typically works, and the failures provide a sense of alternative outcomes that are possible given the same underlying topology of pairwise outcomes. Unpredicted outcomes may occur due to several mechanisms, which are discussed at the last part of the manuscript.

**Figure 3.**
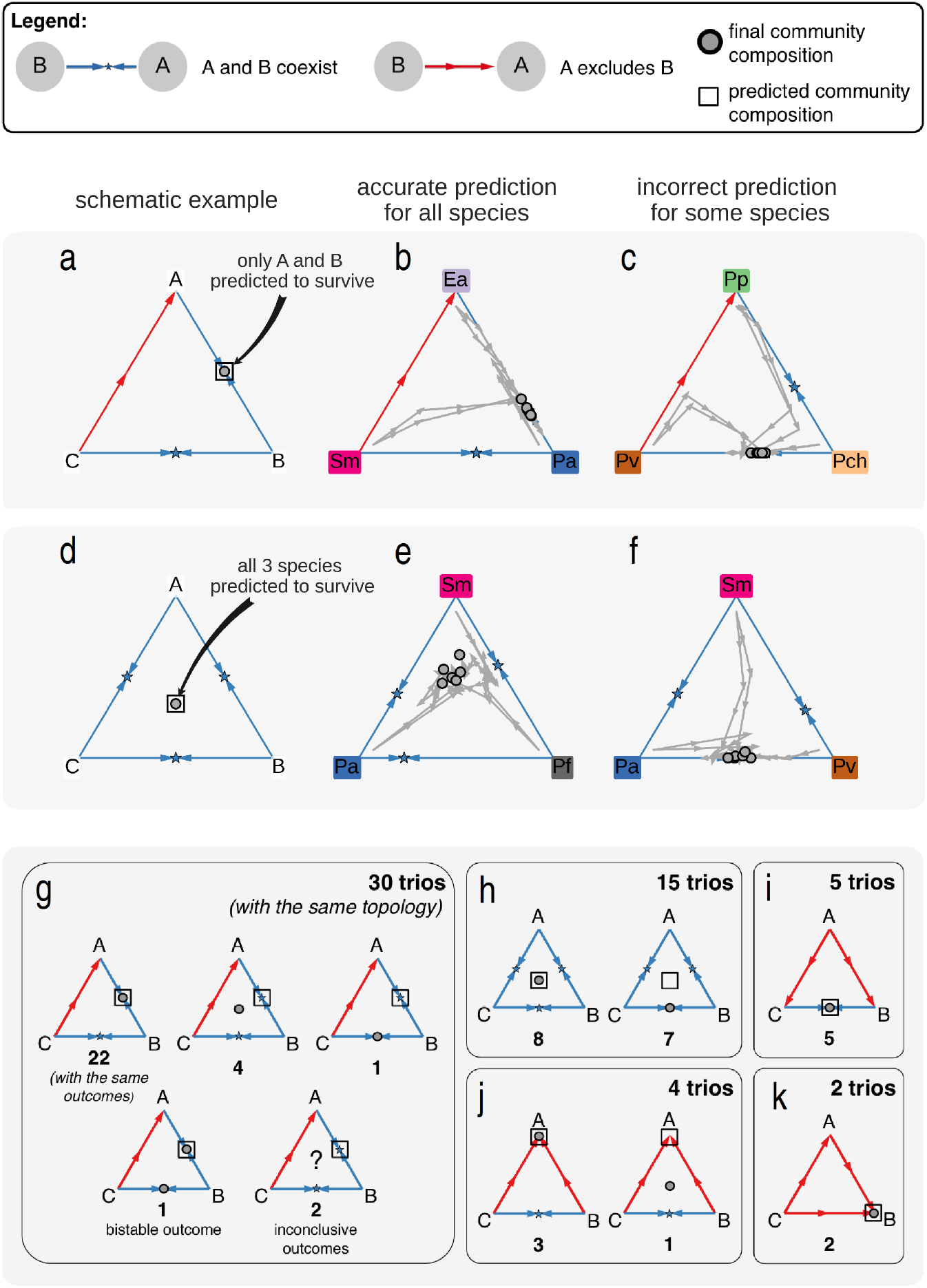
Trio competitions typically resulted in a unique outcome. Changes in species fraction were measured over time for several trio competitions. (a-c) Trios involving 2 coexisting pairs and 1 pair where competitive exclusion occurs. In these plots, each triangle is a simplex denoting the fractions of the three competing species. The simplex vertices correspond to a community composed solely of a single species, whereas edges correspond to a 2-species mixture. The edges thus denote the outcomes of pair competitions, which were performed separately. Trajectories begin at different initial compositions, and connect the species fractions measured at the end of each growth cycle. Dots mark the final community compositions. (a) schematic example, showing that only species A and B are predicted to coexist for this pattern of pairwise outcomes. (b) example of a trio competition which resulted in the predicted outcome. (c) An example of an unpredicted outcome. (d-f) Similar to a-c, but for trios where all species coexist in pairs. (g-k) All trio layouts and outcomes, grouped by the topology of the pairwise outcomes network. Dots denote the final community composition (not exact species fractions, but rather species survivals). One trio displayed bistability, which is indicated by two dots representing the two possible outcomes. Two trios displayed inconsistent results with high variability between replicates, which is indicated by a question mark.

Another frequent topology was coexistence between all three species pairs (15/56 = 27%), in which case none of the species is predicted to be excluded in the trio competition (Fig. 3d). Such trio competitions resulted either in coexistence of all three species, as predicted by our assembly rule (Fig. 3e), or in the exclusion of one of the species (Fig. 3f). Overall, 5 different trio layouts, and 11 competitive outcomes have been observed (Fig. 3g-k). Notably, all observed trio outcomes across all topologies can be generated from simple pairwise interactions, including the outcomes which were not correctly predicted by our assembly rule^24^. An incorrect prediction of our simple assembly rule is therefore not necessarily caused by higher-order interactions.

Overall, survival in three-species competitions was well predicted by pairwise outcomes. The assembly rule predicted species survival across all the three-way competitions with an 89.5% accuracy (Fig. 4a), where accuracy is defined as the fraction of species whose survival was correctly predicted. To get a sense of how the observed accuracy compares to the accuracy attainable when pairwise outcomes are not known, as a null model, we considered the case where the only information available is the average probability that a species will survive in a trio competition (note that this probability is not assumed to be available in our simple assembly rule). Using this information, trio outcomes could only be predicted with a 72% accuracy (Fig. 4a, **Methods**). We further compared the observed accuracy to the accuracy expected when species interact solely in a pairwise manner, according to the gLV equations with a random interaction matrix (**Methods**). We found that the observed accuracy is consistent with the accuracy obtained in simulations of competitions that parallel our experimental setup (p=0.29, Fig. 4b). Survival of species in pairwise competition is therefore surprisingly effective in predicting survival when species undergo trio competition.

**Figure 4.**
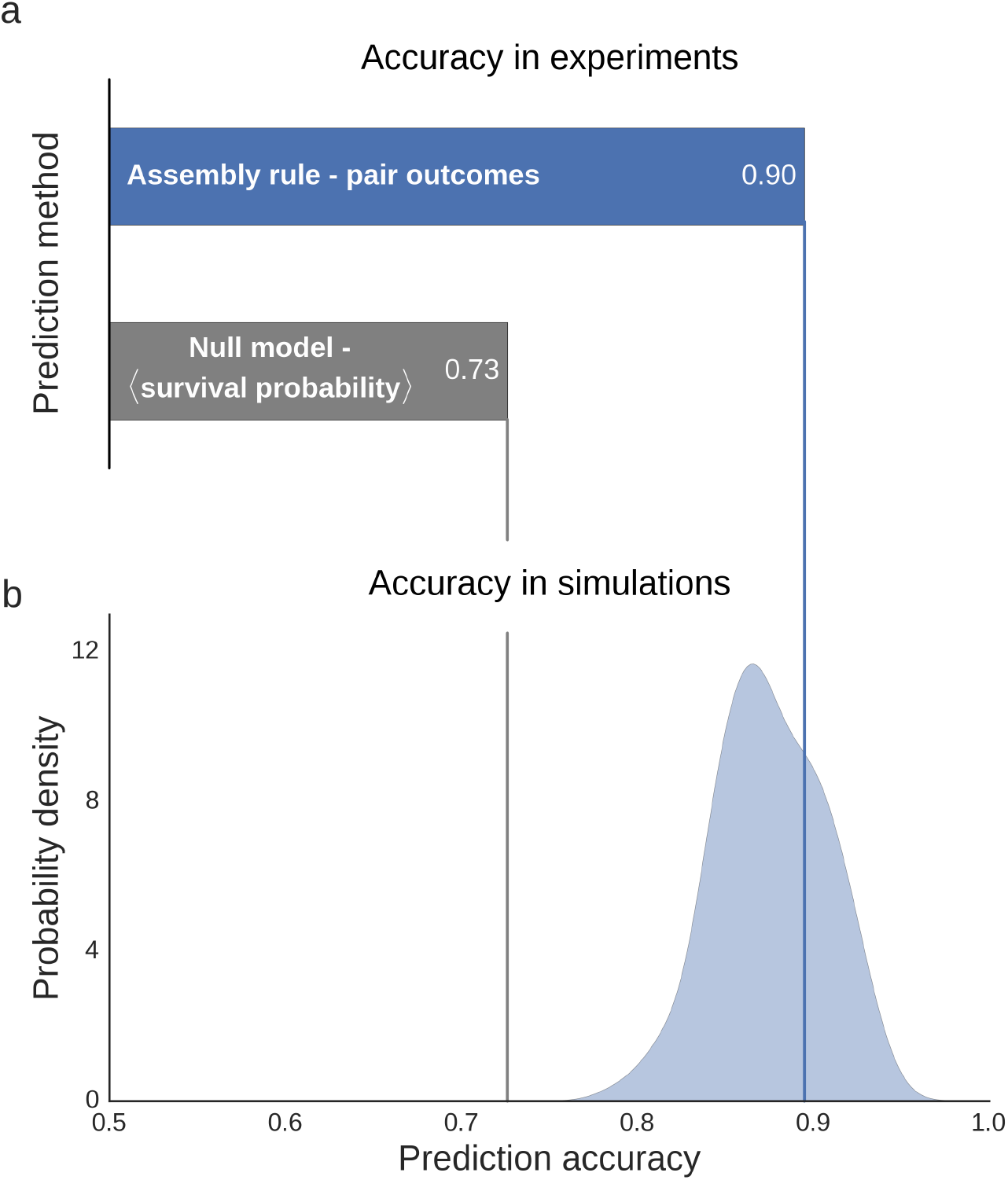
Survival in trio competitions is well predicted by pairwise outcomes. (a) Prediction accuracy of the assembly rule and the null model, where predictions are made solely based on the average probability that species survive in trio competitions. (b) The distribution of accuracies of prediction made using the assembly rule from gLV simulations which mirror our experimental design. The experimentally observed accuracy is consistent with those found in the simulations.

Nonetheless, there are exceptional cases where qualitative pairwise outcomes are not sufficient to predict competitive outcomes of trio competitions. Accounting for such unexpected trio outcomes may improve prediction accuracy for competitions involving a larger set of species. We encode unexpected trio outcomes by creating effective modified pairwise outcomes, which replace the original outcomes in the presence of an additional species. For example, competitive exclusion will be modified to an effective coexistence when two species coexist in the presence of a third species despite one of them being excluded from the pair competition. The effective, modified outcomes can be used to make predictions using the assembly rule as before (**Methods, Fig. S1**). By accounting for unexpected trio outcomes, the assembly rule extends our intuition, and predicts community structure in the presence of potentially complex interactions.

The ability of the assembly rule to predict the outcomes of more diverse competitions was assessed by measuring survival in competitions between all seven-species combinations, as well as the full set of eight species (Fig. 5a). Using only the pairwise outcomes, survival in these competitions could only be predicted with an accuracy of 62.5%, which is barely higher than the 61% accuracy obtained when using only the average probability that a species will survive these competitions (Fig. 5b). A considerably improved prediction accuracy of 86% was achieved by incorporating information regarding the trio outcomes (Fig. 5b). As in the trio competitions, the observed accuracies are consistent with those obtained in gLV simulations that parallel the experimental setup, both when predicting using pairwise outcomes alone (p=0.53), or in combination with trio outcomes (p=0.21, Fig. 5c).

**Figure 5.**
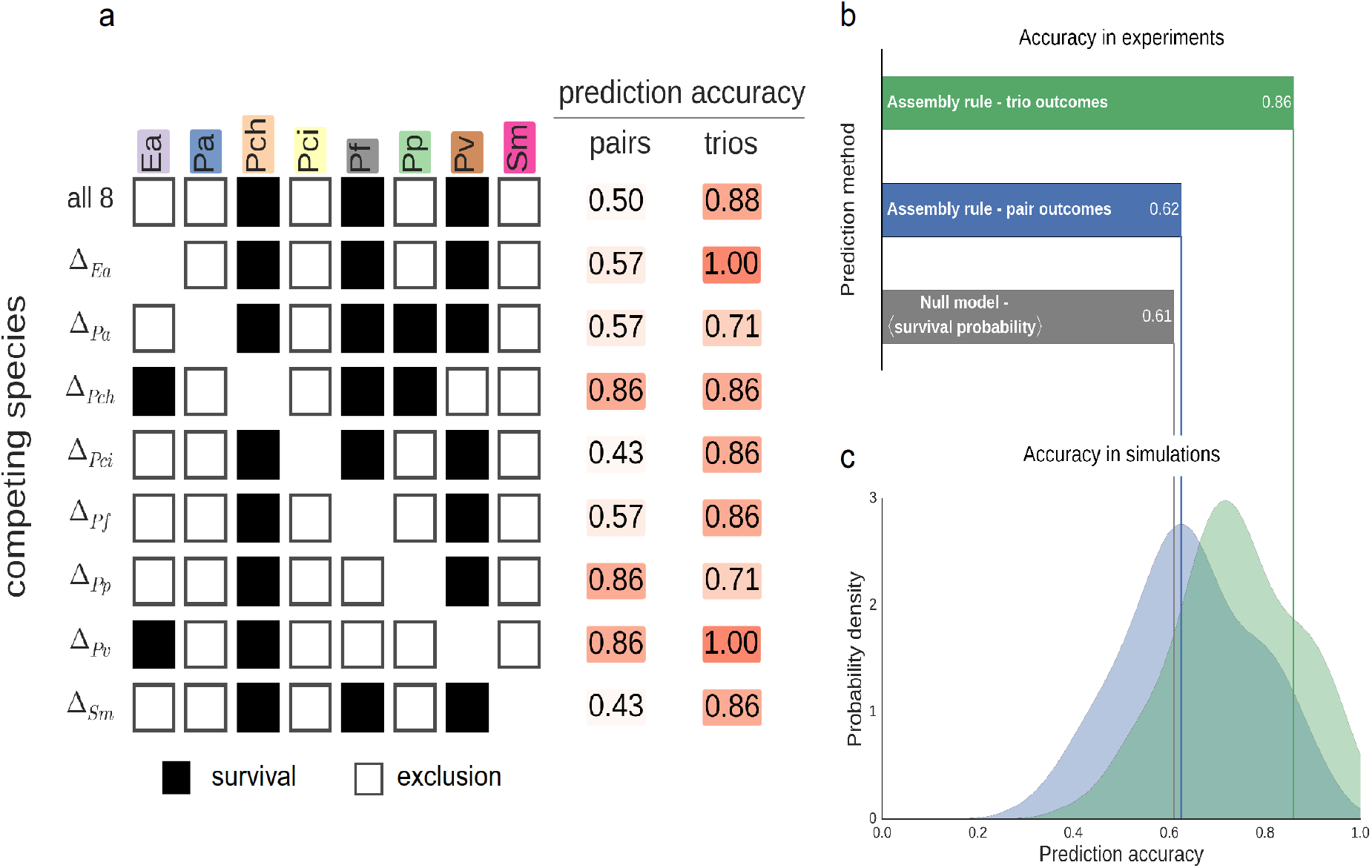
Predicting survival in more diverse competition required incorporating the outcomes of the trio competitions. (a) Species survival when competing all 8 species, and all sets of 7 species. Bright and pale colors indicate survival and extinction, respectively. Survival is predicted either using only pair outcomes, or using both pair and trio outcomes. (b) Prediction accuracy of the null model and the assembly rule, using either pair outcomes only, or pair and trio outcomes (c) The distribution of accuracies of prediction made using the assembly rule from gLV simulations which mirror our experimental design. In these simulations, predictions were made using either pair outcomes only, or pair and trio outcomes. In both cases, the experimentally observed accuracies are consistent with those found in the simulations.

Our assembly rule makes predictions that match our intuition, but there are several conditions under which these predictions may be inaccurate. First, community structure can be influenced by initial species abundances^27^, as has recently been demonstrated in pairwise competitions between bacteria of the genus *Streptomyces*^28^. Our assembly rule may be able to correctly predict the existence of multiple stable states, as it identifies all putative sets of coexisting, non-invasible species in a given species combination. However, we did not have sufficient data to evaluate the rule's accuracy in such cases, as multistability was observed in only one of all our competition experiments.

Complex ecological dynamics, such as oscillations and chaos, can also have a significant impact on species survival^29,30^, making it difficult to predict the community structure. These dynamics can occur even in simple communities containing only a few interacting species. For example, oscillatory dynamics occur in gLV models of competition between as few as three species^24^, and have been experimentally observed in a cross-protection mutualism between a pair of bacterial strains^31^. In contrast, our competitions predominantly resulted in a unique and stable final community. This occurred despite the fact that we observed complex inter-species interactions involving interference competition and facilitation (**Fig S4**). These results indicate that complex ecological dynamics may in fact be rare, though it remains to be seen whether they become more prevalent in more diverse assemblages. Relatedly, prediction is challenging in the presence of competitive cycles (e.g. “Rock-Paper-Scissors” interactions), which often lead to oscillatory dynamics, and are thought to increase species survival and community diversity^32,33^. Such non-hierarchical relationships are absent from our competitive network, and thus their effect cannot be evaluated here.

In the absence of multistability or complex dynamics, our approach may still fail when competitive outcomes do not provide sufficient information regarding the interspecies interactions. This could be due to higher-order interactions, which only manifest in the presence of additional species, or because only qualitative information regarding survival is utilized. The observed accuracy of the assembly rule was consistent with the one found in gLV simulations, but this does not necessarily indicate that our species interact in a linear, pairwise fashion. In fact, fitting the gLV model directly to our pairwise data does not improve predictability (**Fig. S6**). Determining whether, in any particular competition, predictions fail due to insufficient information regarding the strength of linear interactions, non-linear interactions, or higher-order interactions will require more detailed measurements.

Controlling and designing microbial communities has numerous important application areas ranging from probiotic therapeutics, to bioremediation and biomanufacturing^5^. The ability to predict what community will be formed by a given set of species is crucial for determining how extinctions and invasions will affect existing communities, and for engineering desired communities. Our results suggest that, when measured in the same environment, community structure can be predicted from the outcomes of competitions between small sets of species, demonstrating the feasibility of a bottom-up approach to understanding and predicting community structure. While these results are encouraging, they were obtained using a small set of closely related species in well-controlled laboratory settings. It remains to be seen to what extent these results hold in other systems and in more natural settings, involving more diverse assemblages which contain additional trophic levels, in the presence of spatial structure, and over evolutionary time scales.

## Methods

### Species and media

The eight soil bacterial species used in this study are *Enterobacter aerogenes* (Ea, ATCC#13048), *Pseudomonas aurantiaca* (Pa, ATCC#33663), *Pseudomonas chlororaphis* (Pch, ATCC#9446), *Pseudomonas citronellolis* (Pci, ATCC#13674), *Pseudomonas fluorescens* (ATCC#13525), *Pseudomonas putida* (ATCC#12633), *Pseudomonas veronii* (ATCC#700474), and *Serratia marcescens* (Sm, ATCC#13880). All species were obtained from ATCC. The base growth media was M9 minimal media^25^, which contained 1X M9 salts (Sigma Aldrich, M6030), 2mM MgSO4, 0.1mM CaCl2, 1X trace metals (Teknova, T1001). For the final growth media, the base media was supplemented with 1.6mM galacturonic acid and 3.3mM serine as carbon sources, which correspond to 10mM of carbon for each pf these substrates. These carbon sources were chosen from a set of carbon sources commonly used to characterize soil microbes (Biolog, EcoPlate) to ensure that each of the eight species survives in monoculture. Nutrient broth (0.3% yeast extract, 0.5% peptone) was used for initial inoculation and growth prior to experiment. Plating was done on 10cm Petri dishes containing 25mL nutrient agar (nutrient broth with 1.5% agar added).

### Competition experiments

Frozen stocks of individual species were streaked out on nutrient agar Petri plates, grown at room temperature for 48hr, and then stored at 4°C for up to 2 weeks. Prior to competition experiments, single colonies were picked and each species was grown separately in 50mL Falcon tubes, first in 5ml nutrient broth for 24hr and next in 5ml of the experimental M9 media for 48hr. During the competition experiments, cultures were grown in Falcon flat-bottom 96-well plates (BD Biosciences), with each well containing a 150µl culture. Plates were incubated at 25°C without shaking, and were covered with a lid and wrapped in Parafilm. For each growth-dilution cycle, the cultures were incubated for 48hr and then serially diluted into fresh growth media by a factor of 1500.

Initial species mixtures were performed by diluting each species separately to an optical density (OD) of 3*10^−4^. Different species were then mixed by volume to the desired composition. This mixture was further diluted to an OD of 10^−4^, from which all competitions were initialized. For each set of competing species, competitions were conducted from all the initial conditions in which each species was present at 5%, except for one more abundant species. For example, for each species pair there were 2 initial conditions with one species at 95% and the other at 5%, whereas for the 8 species competition there were 8 initial conditions each with a different species at 65% and the rest at 5%. For a few species pairs (Fig. 2a-b), we conducted additional competitions starting at more initial conditions. All experiments were done in duplicate.

### Measurement of cell density and species fractions

Cell densities were assessed by measuring optical density at 600nm using a Varioskan Flash plate reader. Relative abundances were measured by plating on nutrient agar plates. Each culture was diluted by a factor between 10^5^ and 10^6^ in phosphate-buffered saline, depending on the culture's OD. For each diluted culture, 75µl were plated onto an agar plate. Colonies were counted after 48h incubation in room temperature. A median number of 85 colonies per plate were counted. To determine species extinction in competition between a given set of species, we combined all replicates and initial conditions from that competition, and classified as extinct any species whose median abundance was less than 1%, which is just above our limit of detection.

### Assembly rule predictions and accuracy

For any group of competing species, predictions were made by considering all possible competitive outcomes (e.g. survival of any single species, any species pair, etc.). Outcomes that were consistent with our assembly rule were those that were predicted to be a possible outcome of the competition (**Fig. S1**). For any given competition, there may be several such feasible outcomes, however a unique outcome was predicted for all our competition experiments.

Pairwise outcomes were modified using trio outcomes as following: Exclusion was replaced with coexistence for pairs that coexisted in the presence of any additional species. Coexistence was replaced with exclusion whenever a species went extinct in a trio competition with two species with which it coexisted when competed in isolation. Only modifications cause by the surviving species, or an invading species were considered. Therefore, a new set of modified pairwise outcomes was generated for each putative set of surviving species being evaluated.

The prediction accuracy was defined as the fraction of species whose survival was correctly predicted. When the assembly rule identified multiple possible outcomes, which occurred only in the gLV simulations, accuracy was averaged over all such feasible outcomes. Additionally, when the competitive outcome depended on the initial condition, accuracy was averaged across all initial conditions.

For reference, we computed the accuracy of predictions made based on the probability that a species will survive a competition between the same number of species. For example, for predicting trio outcomes, we used the proportion of species that survived, averaged across all trio competitions. Using this information, the highest accuracy would be achieved by predicting that all species survive in all competitions, if the average survival probability is > 0.5, and predicting that all species go extinct otherwise.

### Simulated competitions

To assess the assembly rule's expected accuracy in a simple case in which species interact in a purely pairwise manner, we simulated competitions using the generalized Lotka-Volterra (gLV) dynamics:

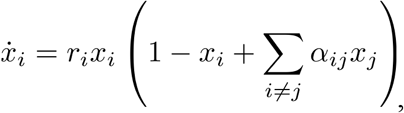

where *x_i_*is the density of species *i*(normalized to its carrying capacity), *r_i_*is the species' intrinsic growth rate, and *α_ij_* is the interaction strength between species *i* and *j*. For each simulation, we created a set of species with random interactions where the *α_ij_* parameters were independently drawn from normal distribution with a mean of 0.6 and a standard deviation of 0.46. Results were insensitive to variations in growth rates, thus they were all set to 1 for simplicity. These parameters recapitulate the proportions of coexistence and competitive exclusion observed in our experiments, and yield a distribution of trio layouts similar to the one (**Fig. S7**). The probability of generating bistable pairs in these simulations is low (~3.7%, corresponding to one bistable pair in a set of eight species), and we further excluded the bistable pairs that were occasionally generated by chance, since we had not observed any such pairs in the experiments.

The accuracy of the assembly rules in gLV systems was estimated by running simulations that parallel our experimental setup: A set of 8 species with random interaction coefficients was generated, and the pairwise outcomes were determined according to their interaction strengths. These outcomes were used to generate predictions for the trio competitions using our assembly rule. Next, all 3-species competitions were simulated with the same set of initial conditions used in the experiments. Finally, the predicted trio outcomes were compared to the simulation outcomes across all trios to determine the prediction accuracy. Thus, a single accuracy value was recorded for each set of 8 simulated species. Similarly, for each simulated 8-species set, the pair and trio outcomes were used to generate predictions for the 7-species and 8-species competitions, and their accuracy was assessed by comparing them to the outcomes of simulated competitions. Prediction accuracy distributions were estimated using Gaussian kernel density estimation from the accuracy values of 100 simulated sets of 8 species.

One-sided P-values evaluating the consistency of the experimentally observed accuracies with the simulation results were defined as the probability that a simulation would yield an accuracy which is at least as high as the experimentally observed one.

### Code availability

An implementation of the assembly rule and the gLV simulations, as well as routines for evaluating the rule's accuracy are freely available online at: https://bitbucket.org/yonatanf/assembly-rule.

## Acknowledgements

We would like to thank A. Perez-Escudero, N. Vega, E. Yurtsev and members of the Gore laboratory for for critical discussions and comments on the manuscript. This work was supported by the DARPA BRICS program, an NIH New Innovator Award (NIH DP2), an NSF CAREER Award, a Sloan Research Fellowship, the Pew Scholars Program and the Allen Investigator Program.

## Author Contributions

J.F. and J.G. designed the study. J.F. and L.H. performed the experiments and analysis. J.F., L.H. and J.G. wrote the manuscript.

## References

1. Berendsen, R. L., Pieterse, C. M. & Bakker, P. A. The rhizosphere microbiome and plant health. Trends Plant Sci. 17, 478–486 (2012).

2. Ezenwa, V. O., Gerardo, N. M., Inouye, D. W., Medina, M. & Xavier, J. B. Animal Behavior and the Microbiome. Science 338, 198–199 (2012).

3. Flint, H. J., Scott, K. P., Louis, P. & Duncan, S. H. The role of the gut microbiota in nutrition and health. Nat. Rev. Gastroenterol. Hepatol. 9, 577–589 (2012).

4. Falkowski, P., Fenchel, T. & Delong, E. The microbial engines that drive Earth’s biogeochemical cycles. Science 320, 1034–9 (2008).

5. Widder, S. et al. Challenges in microbial ecology: building predictive understanding of community function and dynamics. ISME J. (2016). doi:10.1038/ismej.2016.45

6. Großkopf, T. & Soyer, O. S. Synthetic microbial communities. Curr. Opin. Microbiol. 18, 72–77 (2014).

7. Fredrickson, J. K. Ecological communities by design. Science 348, 1425–1427 (2015).

8. Faust, K. & Raes, J. Microbial interactions: from networks to models. Nat. Rev. Microbiol. 10, 538–550 (2012).

9. Bucci, V. & Xavier, J. B. Towards Predictive Models of the Human Gut Microbiome. J. Mol. Biol. 426, 3907–3916 (2014).

10. Berry, D. & Widder, S. Deciphering microbial interactions and detecting keystone species with co-occurrence networks. Front Microbiol 5, 10–3389 (2014).

11. Carrara, F., Giometto, A., Seymour, M., Rinaldo, A. & Altermatt, F. Inferring species interactions in ecological communities: a comparison of methods at different levels of complexity. Methods Ecol. Evol. n/a-n/a (2015). doi:10.1111/2041-210X.12363

12. Billick, I. & Case, T. J. Higher Order Interactions in Ecological Communities: What Are They and How Can They be Detected? Ecology 75, 1530–1543 (1994).

13. Momeni, B. & Shou, W. The validity of pairwise models in predicting community dynamics. bioRxiv 60988 (2016). doi:10.1101/060988

14. Wootton, J. T. The nature and consequences of indirect effects in ecological communities. Annu. Rev. Ecol. Syst. 443–466 (1994).

15. Kelsic, E. D., Zhao, J., Vetsigian, K. & Kishony, R. Counteraction of antibiotic production and degradation stabilizes microbial communities. Nature 521, 516–519 (2015).

16. Stein, R. R. et al. Ecological Modeling from Time-Series Inference: Insight into Dynamics and Stability of Intestinal Microbiota. PLOS Comput Biol 9, e1003388 (2013).

17. Bucci, V. et al. MDSINE: Microbial Dynamical Systems INference Engine for microbiome time-series analyses. Genome Biol. 17, 121 (2016).

18. Vandermeer, J. H. The competitive structure of communities: an experimental approach with protozoa. Ecology 362–371 (1969).

19. Dormann, C. F. & Roxburgh, S. H. Experimental evidence rejects pairwise modelling approach to coexistence in plant communities. Proc. R. Soc. Lond. B Biol. Sci. 272, 1279–1285 (2005).

20. Mounier, J. et al. Microbial Interactions within a Cheese Microbial Community. Appl. Environ. Microbiol. 74, 172–181 (2008).

21. Fisher, C. K. & Mehta, P. Identifying Keystone Species in the Human Gut Microbiome from Metagenomic Timeseries Using Sparse Linear Regression. PLOS ONE 9, e102451 (2014).

22. Marino, S., Baxter, N. T., Huffnagle, G. B., Petrosino, J. F. & Schloss, P. D. Mathematical modeling of primary succession of murine intestinal microbiota. Proc. Natl. Acad. Sci. 111, 439–444 (2014).

23. Case, T. J. & Bender, E. A. Testing for Higher Order Interactions. Am. Nat. 118, 920–929 (1981).

24. Zeeman, M. L. Hopf bifurcations in competitive three-dimensional Lotka-Volterra systems. Dyn. Stab. Syst. 8, 189–216 (1993).

25. Celiker, H. & Gore, J. Clustering in community structure across replicate ecosystems following a long-term bacterial evolution experiment. Nat. Commun. 5, (2014).

26. Concepción-Acevedo, J., Weiss, H. N., Chaudhry, W. N. & Levin, B. R. Malthusian Parameters as Estimators of the Fitness of Microbes: A Cautionary Tale about the Low Side of High Throughput. PLOS ONE 10, e0126915 (2015).

27. Fukami, T. Historical Contingency in Community Assembly: Integrating Niches, Species Pools, and Priority Effects. Annu. Rev. Ecol. Evol. Syst. 46, 1–23 (2015).

28. Wright, E. S. & Vetsigian, K. H. Inhibitory interactions promote frequent bistability among competing bacteria. Nat. Commun. 7, 11274 (2016).

29. Armstrong, R. A. & McGehee, R. Competitive Exclusion. Am. Nat. 115, 151–170 (1980).

30. Huisman, J. & Weissing, F. Biodiversity of plankton by species oscillations and chaos. Nature 402, 407 (1999).

31. Yurtsev, E. A., Conwill, A. & Gore, J. Oscillatory dynamics in a bacterial cross-protection mutualism. Proc. Natl. Acad. Sci. 113, 6236–6241 (2016).

32. Kerr, B., Riley, M. A., Feldman, M. W. & Bohannan, B. J. Local dispersal promotes biodiversity in a real-life game of rock-paper-scissors. Nature 418, 171–174 (2002).

33. Allesina, S. & Levine, J. A. Competitive network theory of species diversity. Proc. Natl. Acad. Sci. 108, 5638 (2011).

